# The single pass membrane protein MRAP2 regulates energy homeostasis by promoting primary cilia localization of the G protein-coupled receptor MC4R

**DOI:** 10.1101/2020.11.13.382325

**Authors:** Adélaïde Bernard, Irene Ojeda Naharros, Florence Bourgain-Guglielmetti, Jordi Ciprin, Xinyu Yue, Sumei Zhang, Erin McDaid, Maxence Nachury, Jeremy F. Reiter, Christian Vaisse

**Affiliations:** Department of Medicine and The Diabetes Center, University of California, San Francisco, California 94143-0540 USA; Department of Ophthalmology, University of California, San Francisco, California 94143-0540 USA; Department of Biochemistry and Biophysics, Cardiovascular Research Institute, University of California, San Francisco, California 94158-2324 USA; Chan Zuckerberg Biohub, San Francisco, CA 94158 USA

## Abstract

The G protein-coupled receptor MC4R (Melanocortin-4 Receptor) and its associated protein MRAP2 (Melanocortin Receptor-Associated Protein 2) are both essential for the regulation of food intake and body weight in humans and mice. MC4R localizes and functions at the neuronal primary cilium, a microtubule-based organelle that senses and relays extracellular signals. Here, we demonstrate that MRAP2 is critical for the ciliary localization and weight-regulating function of MC4R. Our data reveal that GPCR localization to primary cilia can require specific accessory proteins that may not be present in heterologous cell systems. Our findings also demonstrate the essential role of neuronal primary cilia localization of MC4R for adequate control of energy homeostasis and the obesity-promoting effect of genetic disruption of this pathway.

## INTRODUCTION

The regulation of food intake and energy expenditure is dependent on the genetic, molecular and cellular integrity of the central melanocortin system, a network of hypothalamic neurons that integrate peripheral information about energy status and regulate long-term energy homeostasis thereby preventing obesity^1^. This system involves the neuropeptides a-MSH and AGRP, produced by independent populations of neurons in the arcuate nucleus which are sensitive to the adipocyte secreted hormone leptin, as well as the receptor for these neuropeptides, the melanocortin-4 receptor (MC4R). MC4R, one of five members of the Gas-coupled Melanocortin receptor family^2^, is found in multiple brain regions but its expression in the para-ventricular nucleus of the hypothalamus (PVN) is both necessary and sufficient for the regulation of food intake and body weight^3,4^. Underscoring the essential role of this receptor in the maintenance of energy homeostasis, heterozygous coding mutations in *MC4R* are the most common cause of monogenic obesity in humans^5–7^ and of the identified common variants influencing body mass index (BMI), the *MC4R* locus has the second strongest association with obesity^8–10^. Null *Mc4r^-/-^* mice recapitulate the obesity phenotypes observed in humans^11^. Consequently, MC4R is a major target for the pharmacotherapy of obesity, yet little is known about the molecular and cellular pathways underlying the maintenance of long-term energy homeostasis by MC4R expressing neurons.

Recently we reported that MC4R localizes and functions at the neuronal primary cilium^12,13^, a cellular organelle that projects from the surface of most mammalian cell types and functions as an antenna to sense extracellular signals^14^. Mutations disrupting ciliary structure or function, cause ciliopathies^15–17^, genetic disorders characterized by pleiotropic clinical features which can include hyperphagia and severe obesity^18^ such as in Alström or in Bardet-Biedl syndrome (BBS). In adult mice, genetic ablation of neuronal primary cilia also causes obesity^19^.

MC4R interacts with the melanocortin receptor-associated protein MRAP2, a member of the MRAP family comprised of single-pass trans-membrane proteins that interact with Melanocortin receptors^20^ and other GPCRs^21,22^. In non-ciliated heterologous cells, MRAP2 binds to MC4R and increases ligand sensitivity, as well as MC4R-mediated generation of cAMP^23–25^. In human, MRAP2 variants were found in patients with obesity^23,26–28^. *Mrap2^-/-^* mice develop severe obesity although they lack the early-onset hyperphagia of *Mc4r^-/-^* mice^23^ questioning the extent to which MRAP2 qualitatively and quantitatively interacts with MC4R *in vivo.* Indeed, MRAP2 has also been suggested to interact with other GPCRs such as the Ghrelin Receptor and the Prokineticin Receptor^21,22^.

Here we find that the central mechanism of MRAP2-associated obesity is the critical role for MRAP2 in localizing MC4R to cilia.

## RESULTS

### MC4R neurons require MRAP2 to regulate energy homeostasis

Germinal deletion of MRAP2, in *Mrap2^-/-^mice,* leads to obesity although to a lesser extent than that observed in *Mc4r^-/-^* mice^23^. To determine to what extent MC4R-expressing neurons require MRAP2 to control energy homeostasis, we deleted MRAP2 specifically from MC4R-expessing cells. We obtained mice bearing an *Mrap2* knockout-first (“tm1a”) allele and confirmed that, as previously reported^20,23,29^, homozygous mutant mice (hereafter referred to as *Mrap2^-/-^*, Figure S1a) were obese and hyperphagic at 12 weeks of age, but did not differ from their wild-type littermates at 4 weeks of age (Figure S1b,c). From these mice, we generated an *Mrap2* floxed allele (“tm1c” allele, hereafter referred to as *Mrap2^fl^*, Figure S1a). *Mrap2^fl/fl^* mice weighed the same as their wild-type littermates (Figure S1d).

To specifically delete MRAP2 in MC4R-expressing neurons, we generated *Mc4r-t2a-Cre^13^ Mrap2^fl/fl^* mice (hereafter referred to as *Mc4r^t2aCre7t2aCre^ Mrap2^fl/fl^). Mc4r^t2aCre7t2aCre^ Mrap2^fl/fl^* mice fed ad libitum regular chow developed early-onset obesity with significantly higher body weight (Figure 1a-f) and fat mass (Figure 1 k-r), associated with hyperphagia (Figure 1s-v). This phenotype was apparent at 4 weeks of age (Figure 1c,e,k,m,o,q,s,u), and was present both in females and in males (Figure 1). Together, these data demonstrate that MRAP2 is essential in MC4R-expressing neurons to regulate food intake and body weight.

**Figure 1:**
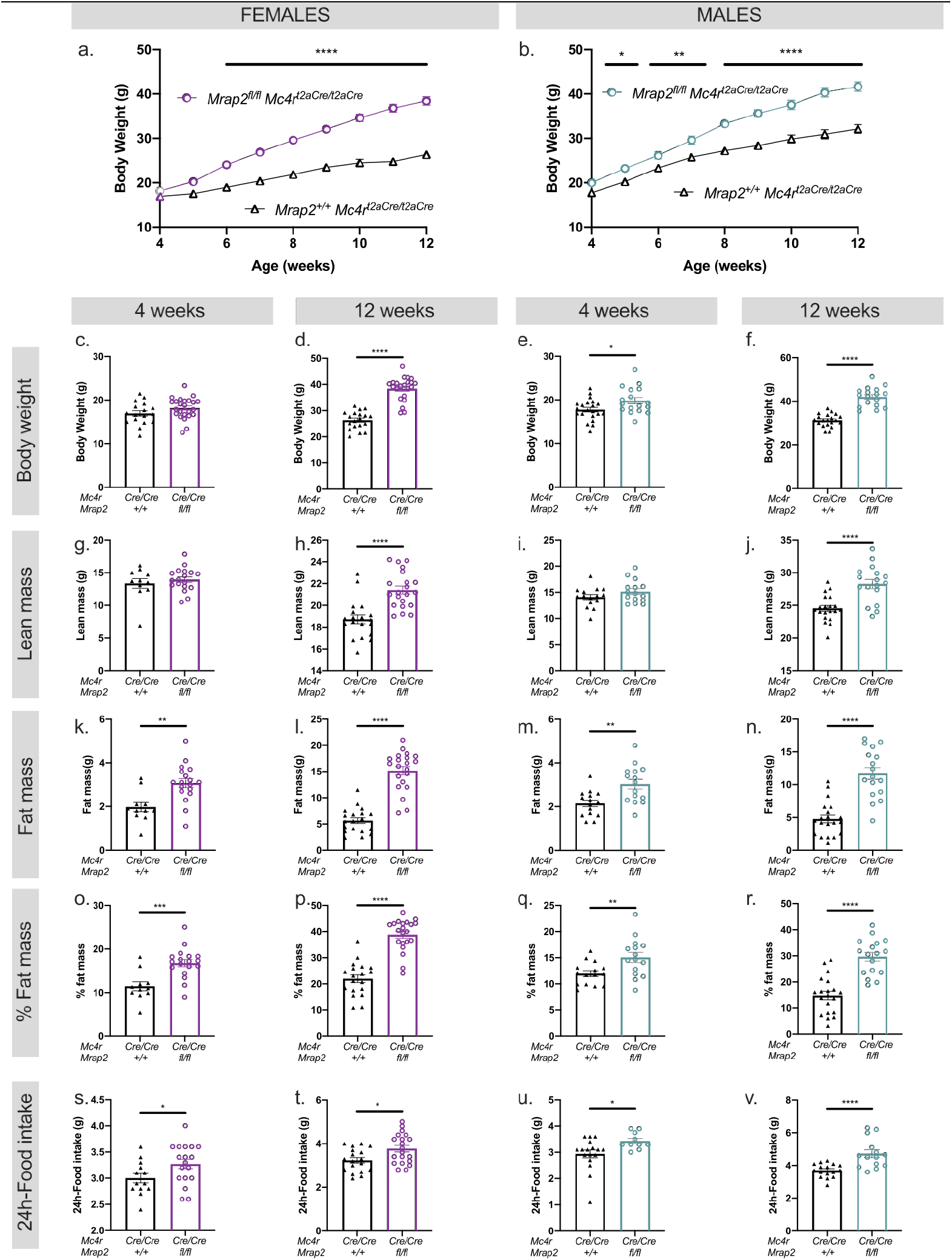
MRAP2 functions in MC4R-expressing cells to regulate food intake and restrain body weight. Body weight curve of *Mc4r^t2aCre/t2aCre^ Mrap2^fl/fl^* vs *Mc4r^t2aCre/t2aCre^ Mrap2^+/+^* female (***a***) and male *(**b**)* mice. Respective body composition at 4 and 12 weeks of *Mc4r^t2aCre/t2aCre^ Mrap2^fl/fl^* vs *Mc4r^t2aCre/t2aCre^ Mrap2^+/+^* females (body weight [***c,d***], lean mass [**g,h**], fat mass [***k,l***], percent fat mass [***o,p***]) and males (body weight [***e,f***|, lean mass *[**ij**],* fat mass [***m,n***], percent fat mass [***q,r***]). 24h-food intake at 4 and 12 weeks of *Mc4r^t2aCre/t2aCre^ Mrap2^fl/fl^* vs *Mc4r^t2aCre/t2aCre^ Mrap2^+/+^* females *(**s,t**)* and males *(**u,v**).* n=11 to n=24 mice per group were used, individual values are displayed. Data are represented as mean ± SEM, *p<0.05, **p<0.01, ***p<0.001, ****p<0.0001, Student’s unpaired t-test (column analysis); mixed-effects model (REML) and Sidak’s multiple comparisons tests (weight curves).

### MRAP2 promotes MC4R localization to the primary cilium in IMCD3 cells

The interaction between MRAP2 and MC4R, as well as the cellular localization of MRAP2 have been previously studied in non-ciliated cells^23–26,30^. Since MC4R localizes to the primary cilium, and MRAP2 has been reported to interact with MC4R *in vitro* ^23^, we tested whether MRAP2 co-localizes with MC4R at the primary cilium following transfection in heterologous cells.

In transiently transfected murine Inner Medullary Collecting Duct (IMCD3) cells, we found that MRAP2 co-localized with the ciliary component, acetylated Tubulin (AcTub, figure S2b), and with MC4R at the primary cilium (Figure 2a, top panel). We tested whether MRAP2 localization is a common feature to the proteins of the MRAP family. We found that a paralog of MRAP2, the Melanocortin Receptor-Associated Protein 1 (MRAP1), an essential accessory factor for the functional expression of the MC2R/ACTH receptor, did not localize to the primary cilium (Figure 2a,b, Figure S2a,b). Ciliary localization is therefore a specific feature of MRAP2.

**Figure 2:**
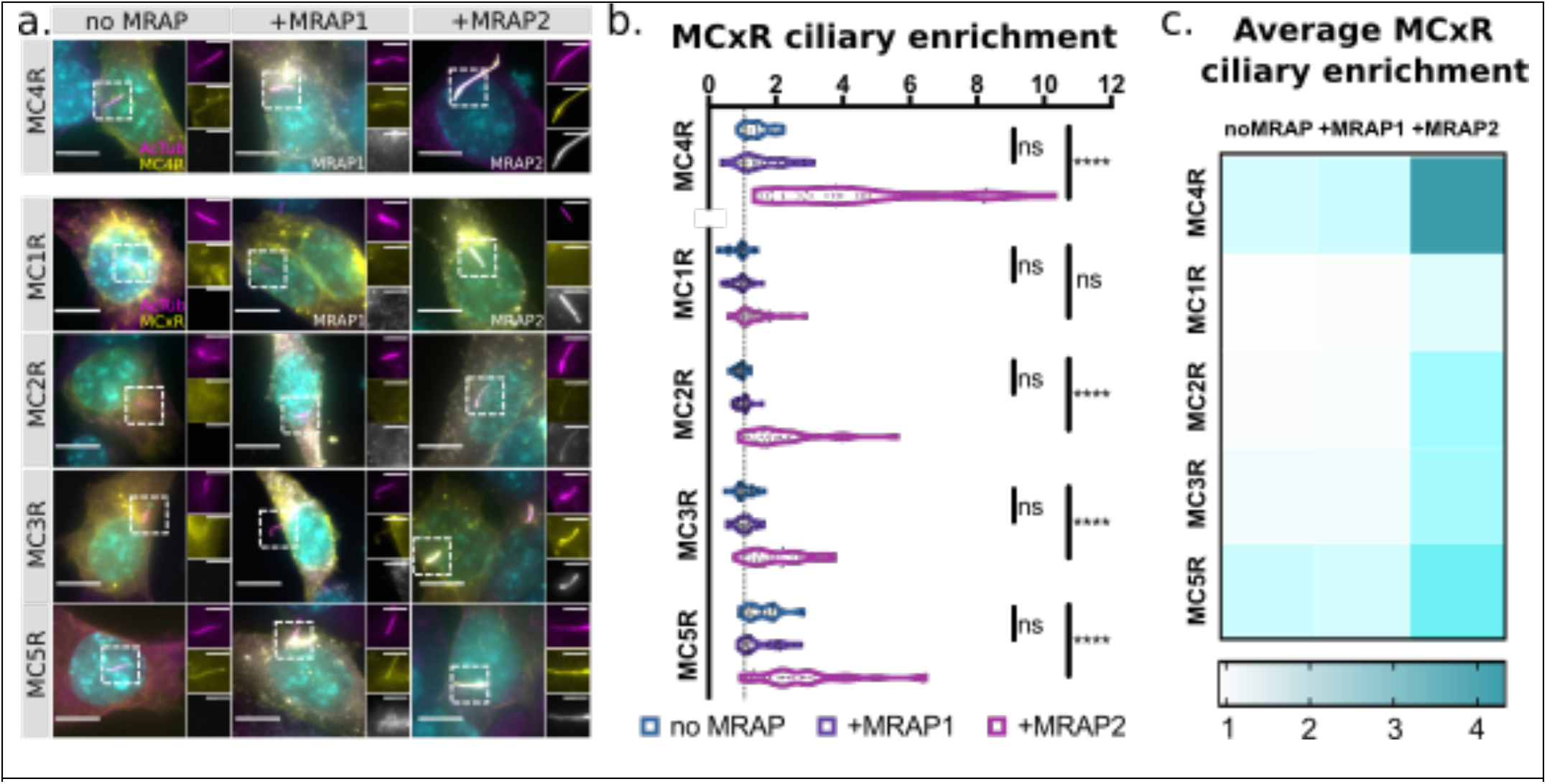
MRAP2 localizes MC4R to the IMCD3 primary cilium. ***a*** Representative widefield micrographs of IMCD3 cells transiently transfected with EGFP-tagged Melanocortin receptors alone (left), or co-transfected with MRAP1-FLAG (center) or MRAP2-FLAG (right). Cells are stained for cilia (AcTub, magenta), EGFP-tagged melanocortin receptors (yellow), MRAP1- or MRAP2-FLAG (white) and nuclei (Hoechst 33342, cyan). MC4R-GFP and MRAP2-FLAG colocalize at the primary cilium (top panel). Scale bars represent 5 μm for low magnification images and 2 μm for the inserts. ***b*** Melanocortin receptor enrichment at the cilium when transfected without MRAP (blue), with MRAP1 (purple) or with MRAP2 (magenta). MRAP2 expression increases ciliary localization of MC2R, MC3R, MC4R and MC5R, but not MC1R. MRAP1 co-expression has no effect on ciliary localization. ***c*** Heatmap displaying mean enrichment at the cilium when MCRs are transfected without MRAP (column 1), with MRAP1 (column 2) or MRAP2 (column 3). MRAP2 highly enriches MC4R localization at the cilium compared to other MCRs. 30-34 ciliated cells per condition were imaged and analyzed. Ciliary and cell body intensity of Melanocortin receptor and MRAP was measured using Fiji. Enrichment at the cilium is expressed as (integrated density at the cilium)/(integrated density in the cell body). Enrichment >1 indicates higher localization of GFP-tagged melanocortin receptor or MRAP at the primary cilium than at the cell body. Data are represented as violin plots. ****p<0.0001, ordinary one-way ANOVA with Sidak’s multiple comparisons test.

We hypothesized that MRAP2 promotes MC4R ciliary localization, and so investigated whether MRAP2 affects MC4R localization. Remarkably, we found that MRAP2 increased MC4R enrichment at the primary cilium, whereas MRAP1 had no effect (Figure 2a,b top panel). Therefore, MRAP2, but not MRAP1, enriches MC4R at primary cilia.

Finally, we assessed the specificity of the interaction between MRAP2 and MC4R by systematically testing the ciliary enrichment of all five Melanocortin receptor family members in the presence or absence of MPRAPs (n=30, Figure 2b,c). Although MRAP2 promoted the ciliary localization of other Melanocortin receptors, the greatest effect was observed on MC4R (Figure 2b,c). MRAP1 did not affect ciliary enrichment of any of the receptors (Figure 2b). Thus, MRAP2 specifically promotes MC4R localization to the primary cilia.

### MRAP2 co-localizes with MC4R at the primary cilia of hypothalamic neurons *in vivo*

To determine whether MRAP2 localizes to the primary cilia *in vivo,* we used a transgenic reporter mouse model in which primary cilia are labelled with GFP *(Arl13b-GFP^tg^*)^31^. MRAP2 sub-cellular localization was assessed by immunofluorescence, which revealed that MRAP2 localizes at the primary cilium in the PVN of these mice (Figure 3a-c). The specificity of the anti-MRAP2 antibody was confirmed by staining hypothalamic sections from *Mrap2^-/-^* mice (Figure S3).

**Figure 3:**
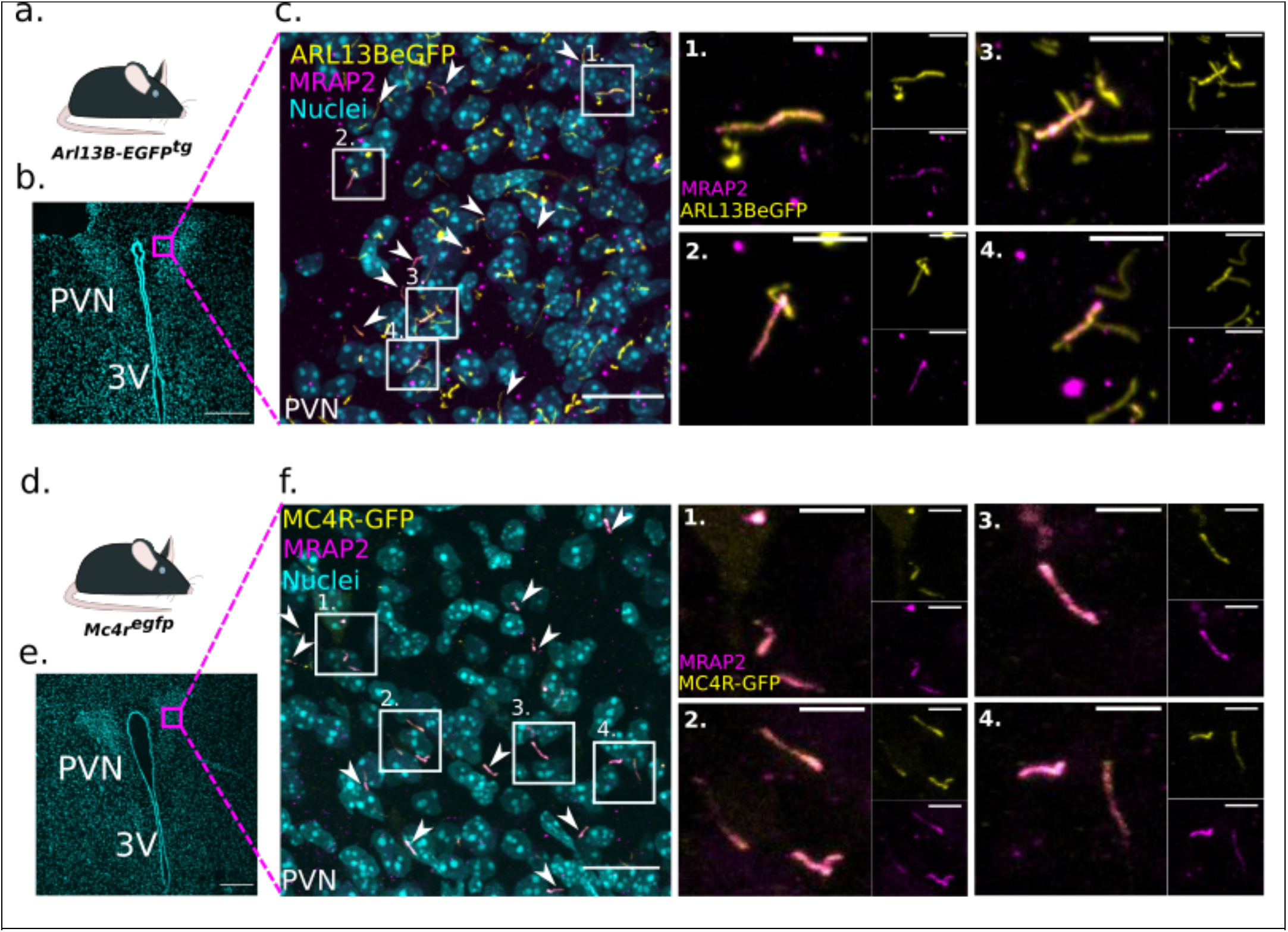
MRAP2 colocalizes with MC4R at the primary cilia *in vivo.* ***a-c*** MRAP2 localizes to primary cilia *in vivo. **a*** Mice expressing a transgene encoding for a ciliary EGFP (Arl13-EGFP) were used in this experiment. ***b*** Representative low magnification image of the PVN, nuclei (Hoescht, cyan). Magenta square indicates higher magnification image depicted in *c*. Scale bar, 200 μm. ***c*** Immunofluorescence image of cilia *(Arl13b-GFP^tg^*, yellow), MRAP2 (magenta) and nuclei (Hoechst, cyan) in the mouse PVN, showing that MRAP2 localizes to primary cilia (arrows). ***d-f*** MRAP2 colocalizes with MC4R *in vivo* (arrows and boxes). ***d*** Mouse line expressing a GFP tag in frame at the C-terminus of the endogenous *Mc4r* locus. ***e*** Representative low magnification image of the PVN, nuclei (Hoescht, cyan). Magenta square indicates higher magnification image depicted in ***f.*** Scale bar, 200 μm. ***f*** Immunofluorescence image of MC4R-GFP (yellow), MRAP2 (Magenta) and nuclei (Hoechst, cyan) in the mouse PVN. Indicated boxed regions are shown to the right at higher magnification. Scale bars, 20 μm for low powered images and 5μm for high powered images. 3V: third ventricle, PVN: paraventricular nucleus of the hypothalamus SCh: suprachiasmatic nucleus.

To further determine whether MRAP2 co-localizes with MC4R at primary cilia, we used a mouse line in which GFP was knocked-in in frame at the C-terminus of the coding sequence of *Mc4r*^12^ (hereafter referred to as *Mc4r^egfp^*) allowing for the assessment of the sub-cellular localization of the endogenous MC4R. In the PVN of *Mc4r^egfp^* mice, most MC4R-GFP localized at primary cilia (Figure S4) and, remarkably, MRAP2 localized exclusively to the cilium of these cells (Figure 3f).

### MC4R localization to primary cilia requires MRAP2

Since MRAP2 enhances MC4R localization at primary cilia *in vitro,* and MC4R co-localizes with MRAP2 at primary cilia *in vivo,* we tested whether loss of MRAP2 compromises MC4R localization at primary cilia *in vivo.* We generated *Mrap2^+/+^ Mc4r^egfp^* and *Mrap2^-/-^ Mc4r^egfp^* mice to compare the ciliary localization of MC4R-GFP in the presence and absence of MRAP2.

In the PVN of *Mrap2^+/+^* mice, MC4R-GFP mainly localized to cilia (Figure 4a). Remarkably, in *Mrap2^-/-^* mice, MC4R-GFP mainly localized to neuronal cell bodies, and was rarely found at primary cilia (Figure 4b). The normalized intensity of MC4R-GFP at cilia (defined by ADCY3 immunostaining) was decreased in *Mrap2* mutants compared to wildtype (Figure 4c, p<0.0001) and MC4R was no longer enriched in cilia in the absence of MRAP2 (Figure 4d, p<0.0001). Thus, MRAP2 is necessary for MC4R enrichment at primary cilia both *in vitro* and *in vivo.*

**Figure 4:**
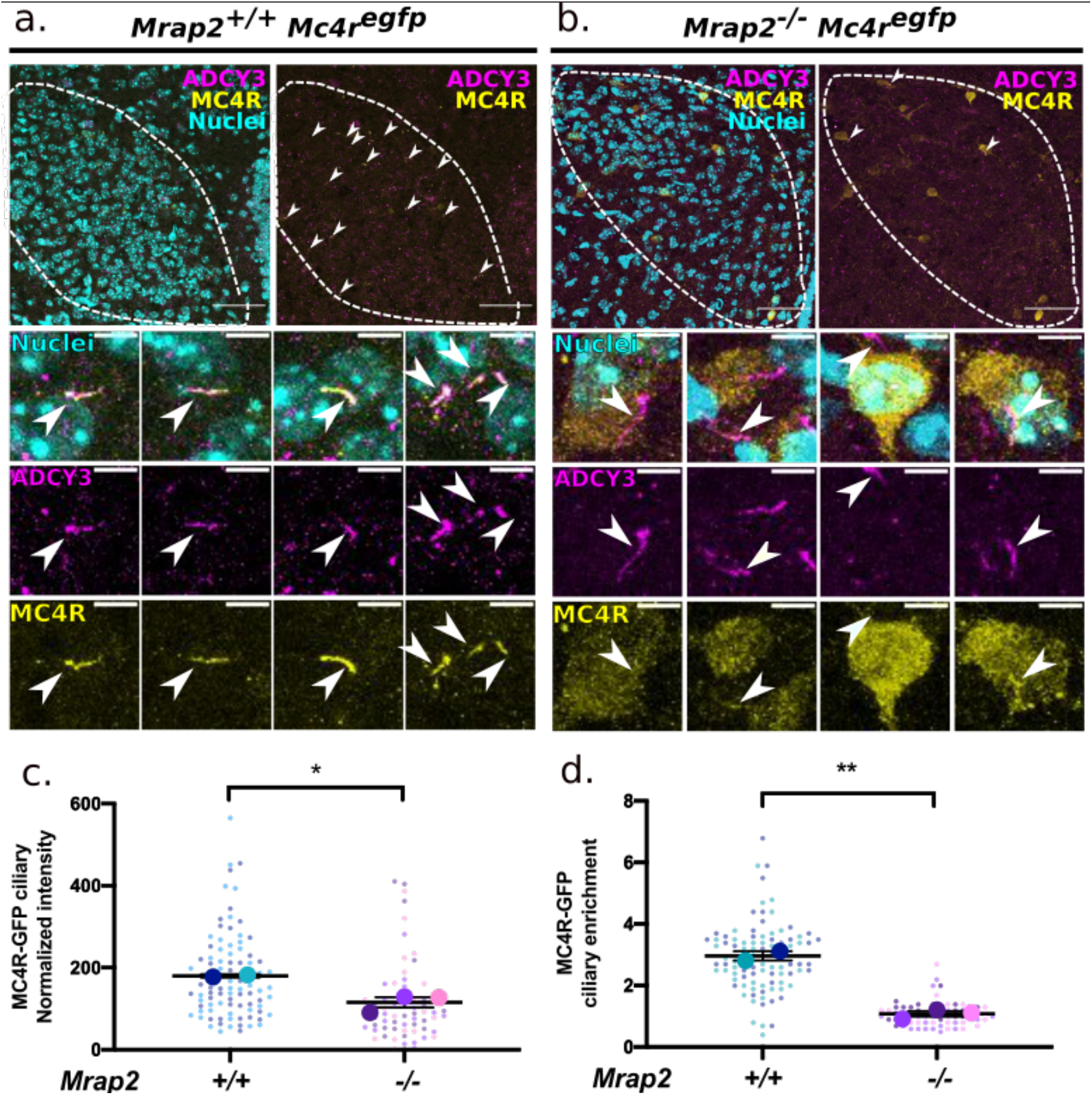
MRAP2 is required for MC4R localization to the primary cilia. ***a,b*** Immunofluorescence images of MC4R-GFP (yellow), cilia (ADCY3, magenta) and nuclei (Hoechst, cyan). Top panels: dashed lines delineate the PVNs. Scale bar, 50 μm. Bottom panels: High magnification images. Scalebar, 5 μm. Arrowheads indicate MC4R-GFP^+^ cilia. ***c*** Quantitation of MC4R-GFP fluorescence intensity at the primary cilium (in arbitrary units). MC4R-GFP ciliary localization is decreased in the absence of MRAP2. ***d*** Quantitation of MC4R-GFP enrichment at the primary cilium as (integrated density at the cilium)/(integrated density in an equal area of the cell body). Enrichment >1 indicates higher localization of MC4R-GFP at the primary cilium than at the cell body. MC4R-GFP localization was quantified from two PVNs per P6 *Mc4r^egfp^ Mrap2^+/+^* (n=2) and *Mc4r^egfp^ Mrap2^-/-^* (n=3) mice. Data are represented as mean ± SEM, *p<0.05, **p<0.01, Student’s t test.

Since whole-body deletion of *Mrap2* could lead to developmental defects potentially accounting for MC4R inability to localize to cilia, we investigated whether acute deletion of *Mrap2* could result in the disruption of MC4R ciliary localization. To test this hypothesis, we removed MRAP2 from the PVN of adult mice. Specifically, we injected *Mrap2^fl/fl^ Mc4r^egfp^* mice unilaterally with an Adeno Associated Virus (AAV) encoding mCherry-IRES-Cre. We predicted that Cre would remove MRAP2 specifically in the infected PVN, and the contralateral uninfected PVN would serve as an internal control (Figure 5b). The brains were harvested 3 weeks following the AAV injections and analyzed. Concordant with our prediction, immunofluorescence staining revealed that, in mCherry-expressing cells, MRAP2 was not apparent at cilia, whereas MRAP2 localized to cilia of the contralateral control PVN (n=4, Figure 5b,c). Cilia, identified by ADCY3 staining, were present in the infected PVN (Figure 5d), confirming that MRAP2 is dispensable for PVN primary cilia maintenance. Remarkably, MC4R localized to cilia in control PVNs, but not in the PVNs from which MRAP2 had been removed (Figure 5d), confirming MRAP2 is essential for MC4R ciliary localization.

**Figure 5:**
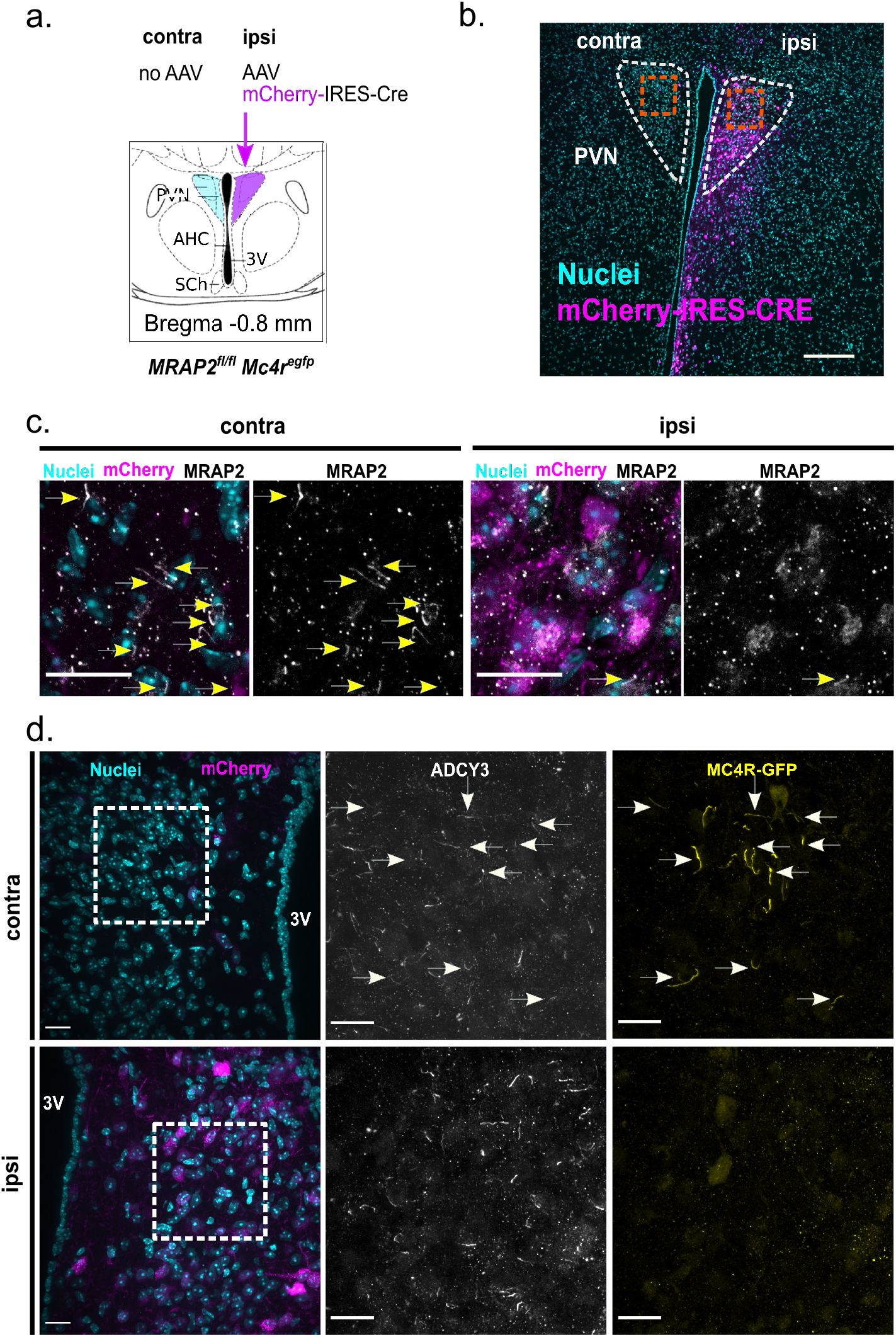
MRAP2 is required in the adult PVN for MC4R ciliary localization. ***a*** Experimental design. *Mrap2^fl/fl^ Mc4r^egfp^* mice were injected unilaterally with AAV encoding mCherry-IRES-Cre (n=4 8-week-old females) and analyzed 3 weeks following injection. ***b*** Representative low magnification image of PVN, mCherry (magenta) and nuclei (Hoescht, cyan). Orange squares indicate higher magnification images depicted in c. Scale bar, 200 μm. ***c*** Higher magnification images of inserts from b. Immunofluorescent staining of MRAP2 (white) of the control contralateral and experimental, ipsilateral PVN with mCherry (magenta) and nuclei (Hoescht, cyan). Arrows indicate MRAP2^+^ cilia, absent from mCherry-expressing cells. Scale bar, 50 μm. ***d*** Immunofluorescent staining of control, contralateral PVN and experimental, ipsilateral PVN for, on the left, mCherry (magenta) and nuclei (Hoescht, cyan), in the middle, ADCY3 (white) and, on the right, MC4R-GFP (yellow). White squares indicate regions imaged for higher magnification images in the middle and right. ADCY3 is not altered by loss of MRAP2 while MC4R-GFP localization to neuronal primary cilia is abrogated by loss of MRAP2. Arrows indicate MC4R^+^ cilia. Scale bars, 20 μm.

To test whether MRAP2 contributes to the ciliary localization of other GPCRs, we assessed the localization of the Somatostatin Receptor 3 (SSTR3) in the presence or absence of MRAP2. SSTR3 is a cilia-localized GPCR which is also expressed in the PVN^32^, including a subset of MRAP2-expressing neurons (Figure S5d,e). Interestingly, the absence of MRAP2 did not affect ciliary localization of SSTR3 (Figure S6). We conclude that MRAP2 is specifically required for MC4R localization at the primary cilium *in vivo.*

## DISCUSSION

Ablating primary cilium by conditionally knocking out *Ift88* in adult mice leads to obesity, either when deleted ubiquitously, specifically in neurons, or only in the PVN, directly implicating PVN cilia in the control of feeding behavior^13,19^.

The composition of the primary ciliary membrane is different from that of the surrounding plasma membrane, as it is enriched for proteins involved in specific forms of signaling^33^. We previously identified MC4R as one of a select subset of GPCRs that localizes to cilia. Moreover, blocking Gs signaling specifically at the primary cilium of MC4R-expressing PVN neurons also leads to obesity^13^, suggesting that not only does MC4R localize to cilia, but it functions there to cue energy homeostasis. Importantly, human obesity-associated *MC4R* mutations affecting its third intracellular loop (a domain previously implicated in the ciliary localization of other GPCRs^34^) impair MC4R ciliary localization without affecting its trafficking to the cell membrane or its ability to couple to G proteins^12^, providing evidence that ciliary localization is critical for MC4R function in humans. Given the functional connection between cilia and MC4R function, we sought insights into regulators of MC4R ciliary localization.

In this study, we demonstrate that a ciliary GPCR accessory protein, MRAP2, restrains feeding by acting in MC4R-expressing neurons to direct its associated GPCR, MC4R, to cilia. As MC4R and MRAP2 are essential for suppressing feeding behavior and MC4R fails to localize to primary cilia in *Mrap2^-/-^* mice, we propose that the failure of MC4R to localize to cilia accounts for the obesity observed in MRAP2-deficient mice and humans.

Previous *in vitro* studies in unciliated cells have reported opposing effects of MRAP2 on MC4R function: one study reported that MRAP2 does not affect MC4R activity^30^, another suggested that MRAP2 may inhibit MC4R activity^24^, and others that MRAP2 increases its response to ligand^23,25^. MRAP2 was also reported to either positively or negatively regulate MC4R activity depending on their relative concentrations^26^. Similarly, MRAP2 has been reported to either decrease^24^ or modestly increase^25^ MC4R cell-surface expression. It will be of interest to repeat these assays in ciliated cells.

The ciliary trafficking of GPCRs such as SSTR3, HTR6 and Rhodopsin share their dependence on interactors such as Tubby family members^32,35–37^. In contrast, the dependence of MC4R on MRAP2 for ciliary trafficking may reveal a more specific requirement. For example, we found that, although MRAP2 and SSTR3 co-localize at primary cilia of some PVN neurons, SSTR3 does not require MRAP2 for ciliary localization. Therefore, MRAP2 may be a ciliary trafficking chaperone for MC4R, rather than a component of the general ciliary trafficking machinery. Ciliary accessory proteins, such as MRAP2, may be a site of regulation for the ciliary localization and function of their associated receptors.

Although MRAP2 is not required generally for ciliary GPCR localization, it can associate with other GPCRs beyond MC4R, including GHSR1a^22^ and PKR1^21^. It will be interesting to assess whether other MRAP2-associated receptors localize to cilia in MRAP2-dependent ways. Association of MRAP2 with receptors other than MC4R could also explain the differences in weight phenotypes caused by removing *Mrap2* globally and specifically in MC4R-expressing neurons. Specifically, we found that inactivating *Mrap2* specifically in MC4R-expressing neurons causes, like *Mc4R* loss-of-function, early onset hyperphagia and obesity, whereas germline inactivation of *Mrap2* caused a milder phenotype, with late onset hyperphagia and obesity^23,29^. Therefore, it will be interesting to assess whether MRAP2 promotes the ciliary function of orexigenic receptors.

The evident role of primary cilia in metabolism^12,18,38–40^ has motivated *in vitro* screens to identify ciliary GPCRs that may control energy homeostasis. Previously described ciliary GPCRs include the Melanin-Concentrating Hormone Receptor 1 (MCHR1)^34^ and the Neuropeptide Y Receptor type 2 (NPY2R)^41,42^. However, the necessity of an accessory protein like MRAP2 for ciliary trafficking of specific GPCRs was not considered, suggesting that these screens may have missed a number of ciliary GPCRs, including MC4R^41,42^. Whether other ciliary GPCRs use accessory proteins for ciliary trafficking is an open question, but is hinted at by studies demonstrating that proteins, such as Rhodopsin, localize robustly to cilia in their native cell types, but less well in heterologous ciliated cells^43^.

Our study revealed that MRAP2 is required for MC4R localization to and function at primary cilia and its resulting function, and is therefore necessary to regulate long-term energy homeostasis. The emerging connection between primary cilia and energy homeostasis^12,18,38–40^ suggests that mutations in other genes necessary for the ciliary localization and function of MC4R will be candidate causes for human obesity.

## MATERIAL AND METHODS

### Cell culture and Transfections

#### Expression plasmids

MC1R-GFP, MC2R-GFP, MC3R-GFP and MC5R-GFP expression constructs were constructed as previously described for MC4R-GFP^44^. Plasmids encoding MRAP1-FLAG and MRAP2-FLAG were obtained from Dr. Hinkle^30^.

#### Ciliary expression of MCRs and MRAPs in cultured cells

IMCD3 cells were transfected using X-tremeGENE^TM^ 9 DNA Transfection Reagent (06365809001, Roche). The transfection reagent was diluted in OptiMEM (Life Techonologies) and incubated at room temperature for 5 min. Then, the mixture was added to the diluted plasmids in a 6:1 ratio (6 μl transfection reagent to 1 ug DNA). 50,000 cells in suspension were added to the transfection mixture after 20 min incubation at room temperature. Transfected cells were switched to starvation media after 24 h and fixed after 16 h. Double plasmid transfections were done by diluting equal mass of each vector.

#### Cell imaging

50,000 cells were seeded for transfection on acid-washed 12 mm #1.5 cover glass (Fisherbrand) in a 24-well plate. Starved cells were fixed in phosphate buffered saline (PBS) containing 4% paraformaldehyde (Electron Microscopy Sciences) for 15 min at room temperature and permeabilized in ice-cold 100% methanol (Fisher Scientific) for 5 min. Cells were then further permeabilized in PBS containing 0.1% Triton X-100 (BP151-500, Thermo Fisher Scientific), 5% normal donkey serum (017000-121, Jackson Immunoresearch Labs), and 3% bovine serum albumin (BP1605-100, Thermo Fisher Scientific) for 30 min. Permeabilized cells were incubated with the specified primary antibodies (mouse monoclonal anti-acetylated tubulin T6793, Sigma-Aldrich; and mouse monoclonal anti-FLAG M2 antibody, F1804, Sigma-Aldrich) for 1 h washed with PBS, and incubated with dye-coupled secondary antibodies (Jackson Immunoresearch Labs) for 30 min. Cells were then washed with PBS, stained with Hoechst DNA dye, and washed with PBS before mounting with Fluoromount G (Electron Microscopy Sciences).

Cells were imaged in a widefield fluorescence DeltaVision microscope (Applied Precision) equipped with a PlanApo 60x/1.40NA objective lens (Olympus), a pco.edge 4.2 sCMOS camera, a solid state illumination module (Insight) and a Quad polycroic (Chroma). Z stacks with 0.2 μm separation between planes were acquired using SoftWoRx. The illumination settings were: 140 μW 390 nm wavelength for 0.15 s to image Hoechst, 222 μW 475 nm wavelength for 0.3 s to image Alexa Fluor 488-stained MCRs, 123 μW 543 nm wavelength for 0.3 s to image Cy3-stained acetylated tubulin and 115 μW 632 nm wavelength for 0.15 s to image Cy5-stained FLAG-labelled MRAPs. Images were flat field-corrected, background subtracted and maximally projected using Fiji. Ciliary intensity measurements were also taken in Fiji.

### *In vivo* experiments

#### Animals

Mice were housed in a barrier facility and maintained on a 12:12 light cycle (on: 0700-1900) at an ambient temperature of 23±2°C and relative humidity 50-70%. Mice were fed with rodent diet 5058 (Lab Diet) and group-housed up to 5. Experiments were performed with weight matched littermates. All animal procedures were approved by the Institutional Animal Care and Use Committee of the University of California, San Francisco.

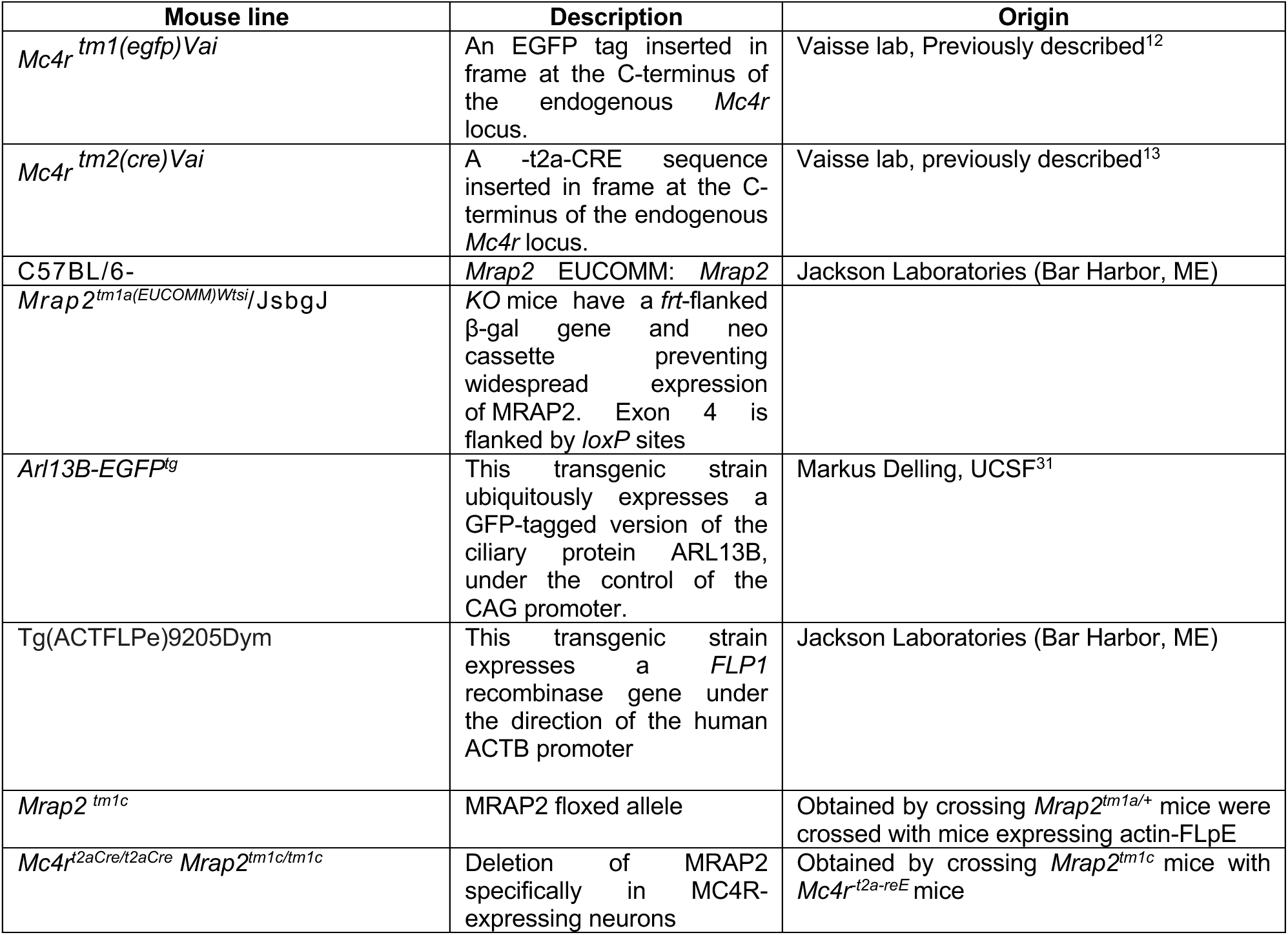

#### EUCOMM MRAP2 mice

EUCOMM MRAP2 knockout-First allele (“tm1a”) mice carry an *frt*-flanked β-gal gene and neo cassette preventing widespread expression of the *Mrap2* gene (EUCOMM tm1a allele or *Mrap2*^-/-^). When mice harboring this allele are crossed into an actin-flip background, the frt-flanked cassette is excised and MRAP2 wild-type function is restored (EUCOMM tm1c allele or *Mrap2^fl/fl^).* After Flip-mediated excision, a loxP-flanked Exon 4 remains, which allows for MC4R cell-specific deletion when crossed to *Mc4r-t2a-CRE* knock-in mice *(Mc4r^t2aCre/t2aCre^ Mrap2^fl/fl^*).

#### Stereotaxic AAV-injection Surgeries

8 weeks old *Mrap2^fl/fl^ Mc4r^egfp^* females (n=4) were injected unilaterally with pAAV-Ef1a-mCherry-IRES-CRE (Addgene, catalog #55632-AAV8).

Animals were anesthetized with an initial flow of 4% isoflurane, maintained under anesthesia using 2% isoflurane and kept at 30-37°C using a custom heating pad. The surgery was performed using aseptic and stereotaxic techniques. Briefly, the animals were put into a stereotaxic frame (KOPF Model 1900, USA), the scalp was opened, the planarity of the skull was adjusted and a hole was drilled (PVN coordinates: AP=-0.8, ML=-0.2, DV=-5.3). A volume of 300 nl was injected at a rate of 0.1 μL/min. Animals were given pre-operative analgesic (buprenorphine, 0.3 mg/kg) and post-operative anti-inflammatory Meloxicam (5 mg/Kg) and allowed to recover at least 10 days during which time they were single-housed and handled frequently. The mice were calorie-restricted at 75% for 2 weeks prior to perfusion.

#### Brain imaging

Mice were perfused trans-cardially with PBS followed by 4% paraformaldehyde fixation solution. Brains were dissected and post-fixed in fixation solution at 4°C overnight, soaked in 30% sucrose solution overnight, embedded in O.C.T. (Tissue-Tek, Sakura Finetek USA, INC., Torrance, CA), frozen, and cut into 20-35 μm coronal sections, then stored at −80°C until staining.

After washing, sections were blocked for 1 h in 50% serum 50% antibody buffer (1.125% NaCl, 0.75% Tris base, 1% BSA, 1.8 % L-Lysine, 0.04% sodium azide), followed by incubation with primary antibody overnight at 4°C. After washing, sections were incubated with secondary antibodies for 1 h at room temperature, washed and stained with Hoechst (1:5000), washed and mounted with Prolong™ Diamond antifade Mountant.

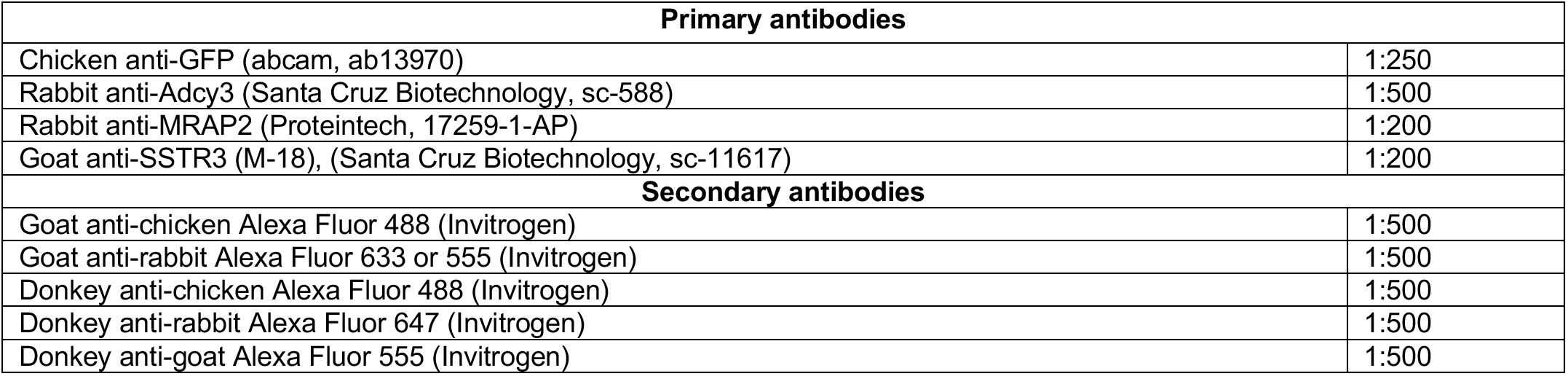

| Primary antibodies |  |
| --- | --- |
| Chicken anti-GFP (abcam, ab13970) | 1:250 |
| Rabbit anti-Adcy3 (Santa Cruz Biotechnology, sc-588) | 1:500 |
| Rabbit anti-MRAP2 (Proteintech, 17259-1-AP) | 1:200 |
| Goat anti-SSTR3 (M-18), (Santa Cruz Biotechnology, sc-11617) | 1:200 |
| Secondary antibodies |  |
| Goat anti-chicken Alexa Fluor 488 (Invitrogen) | 1:500 |
| Goat anti-rabbit Alexa Fluor 633 or 555 (Invitrogen) | 1:500 |
| Donkey anti-chicken Alexa Fluor 488 (Invitrogen) | 1:500 |
| Donkey anti-rabbit Alexa Fluor 647 (Invitrogen) | 1:500 |
| Donkey anti-goat Alexa Fluor 555 (Invitrogen) | 1:500 |

In figure 5, the immunofluorescence stainings were performed on brain sections from mice that were calorie-restricted for a week at 75% of baseline food intake on regular chow and fasted for 24 h prior to perfusion. In figure S3 (validation of MRAP2 antibody), the staining for MRAP2 and GFP was performed on brain sections from P6 mice expressing MC4R-GFP, either wild type or knock out for MRAP2 (MRAP2 1a/1a).

#### Microscopy

| Microscope | Figure |
| --- | --- |
| Leica SP5 (Z-stack, 0.5 μm steps) | Assessment of MC4R-GFP at the cilium of MRAP2 wt vs KO |
| Leica SP8 with resonant scanner | Colocalization of MC4R-GFP and MRAP2 in P6 mice |
| Nikon W1 wide field-of-view spinning disk confocal with Andor Zyla sCMOS camera, (Z-stack, 0.26 μm steps) | SSTR3 imaging, MRAP2 IF in MC4R-GFP adults |

#### Image processing

Images were processed with Fiji. Maximal intensity Z projections are from at least 20 slices over 15-20 μm. Quantified slices were matched for the number of slices projected and settings.

#### Quantification of ciliary localization in cultured cells and hypothalamus sections

Matched Z-stack maximum projections were analyzed in Fiji. Relative ciliary enrichment was calculated as follows: each primary cilium was manually defined by a segmented line following ADCY3+ signal, and the pixel intensity in other channels was measured in that defined area (integrated density, IntDen). Ciliary intensity of MC4R-GFP was then calculated as the IntDen of MC4R-GFP in the cilium, subtracting adjacent background (measured as IntDen of same defined area nearby the cilium). To calculate relative cilia enrichment, the IntDen (cilium) was divided by the Intden (cell body), measured in the closest cell body (as defined by the presence of a Hoechst positive nucleus). Enrichment >1 therefore indicates higher localization of the receptor at the primary cilium compared to cell body.

#### Mouse Metabolism Studies

For experiments presented in Figure 1, mice were weaned at 4 weeks of age, single-housed and their food intake was measured manually every 24 h for 4 consecutive days and averaged. Food intake data was excluded if the mouse lost a significant amount of weight because of single housing stress. The animals were then housed in groups of five, and their weight was measured weekly until 12 weeks of age. Food intake was again assessed as described at 12 weeks of age. Body composition was assessed by EchoMRI™ at 4 and 12 weeks of age.

#### Statistics

Sample sizes were chosen based upon the estimated effect size drawn from previous publications and from the performed experiments. Data distribution were assumed to be normal, but this was not formally tested. All tests used are indicated in the figures. We analyzed all data using Prism 7.0 (GraphPad Software). (^ns^p>0.05; *P≤0.05; **P≤0.01; ***P≤0.001; ***P≤0.0001).

#### Data availability

The data that support the findings of this study are available from the corresponding author upon request.

## Notes

**Sources of support:** This research was supported by NIH (R01AR05439, R01GM095941) to JFR, NIH R01 DK 60450 to CV, NIH R01DK106404 to JFR and CV, NIH GM089933 to MVN, a Sandler Integrative Research Award to CV and JFR, a Stein Innovation Award from Research to Prevent Blindness to MVN, a grant from the Swiss National Science Foundation (P2ZHP3_178137) to ION, as well as NEI core grant EY002162 and the UCSF NORC (NIH P30DK098722).

### Competing Interest Statement

The authors have declared no competing interest.

