## Supplemental data for "The single pass membrane protein MRAP2 regulates energy homeostasis by promoting primary cilia localization of the G protein-coupled receptor MC4R"

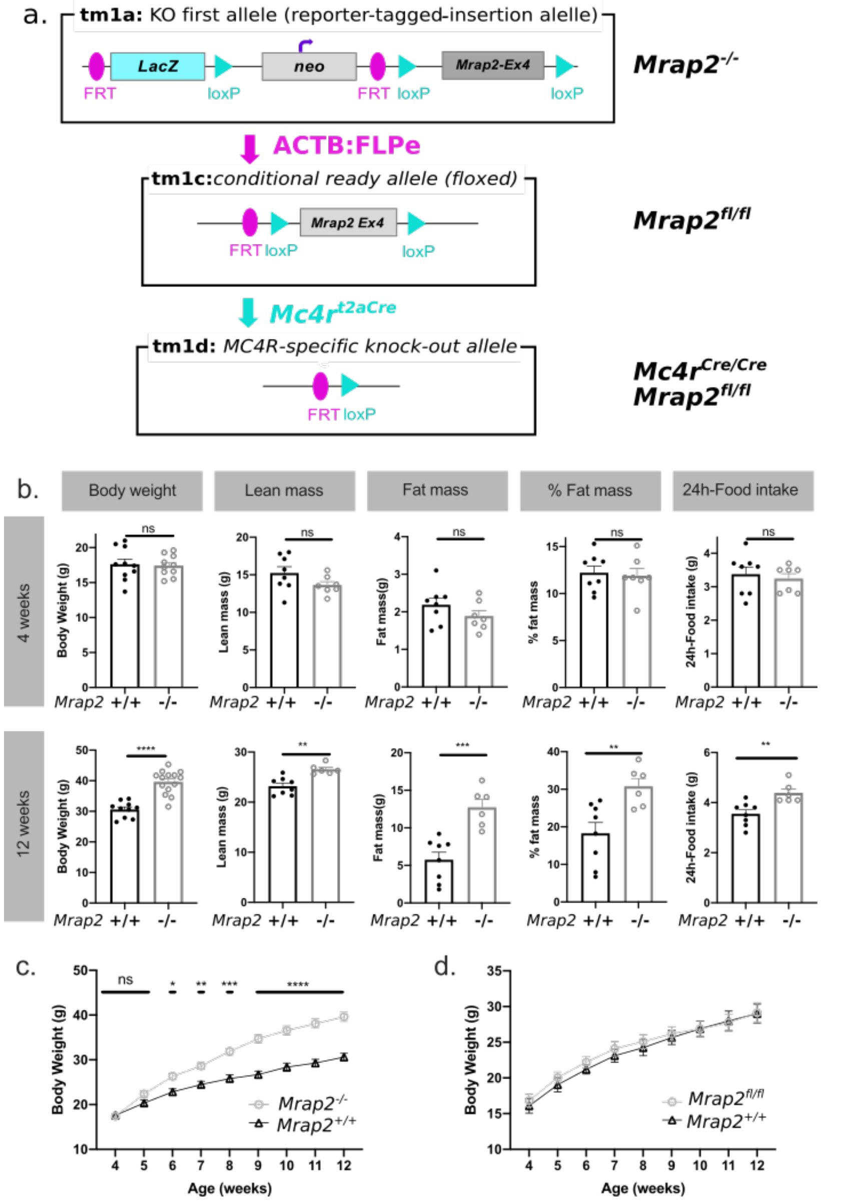


**Figure S1: Phenotypical characterization of mice bearing EUCOMM MRAP2 tm1a and tm1c alleles.** ***a*** EUCOMM allele nomenclature and specific crosses. EUCOMM *Mrap2* knockout-First allele ("tm1a") mice carry an *frt*-flanked β-gal gene and neo cassette preventing widespread expression of the *Mrap2* gene (EUCOMM tm1a allele or *Mrap2*$^{-/-}$). When mice harboring this allele are crossed into an actin-flip background (ACTB:FLPe), the frt-flanked cassette is excised and *Mrap2* wild-type function is restored (EUCOMM tm1c allele or *Mrap2*$^{fl/fl}$). After Flip-mediated excision, a loxP-flanked Exon 4 remains, which allows for *Mc4r* cell-specific deletion when crossed to *Mc4r-t2a-CRE* knock-in mice (*Mc4r*$^{t2aCre/t2aCre}$ *Mrap2*$^{fl/fl}$). ***b*** Body composition and 24h-food intake at 4 and 12 weeks of age (top and bottom panel respectively) of male mice homozygous for the EUCOMM *Mrap2*$^{tm1a}$ allele (*Mrap2* whole body knockout, *Mrap2*$^{-/-}$, n=14) compared to their wildtype littermates (*Mrap2*$^{+/+}$, n=10). ***c*** Body weight curve of *Mrap2*$^{-/-}$ (n=14) compared to wildtype *Mrap2*$^{+/+}$ littermates (n=10). ***d*** Body weight curve of male mice homozygous for the EUCOMM *Mrap2*$^{tm1c}$ allele (*Mrap2* floxed allele, *Mrap2*$^{fl/fl}$, n=7), compared to their wildtype littermates (*Mrap2*$^{+/+}$, n=10). Data are represented as mean ± SEM, *p<0.05, **p<0.01, ***p<0.001, ****p<0.0001, Student's unpaired t-test (column analysis); Mixed-effects model (REML) and Sidak's multiple comparisons tests (weight curves).

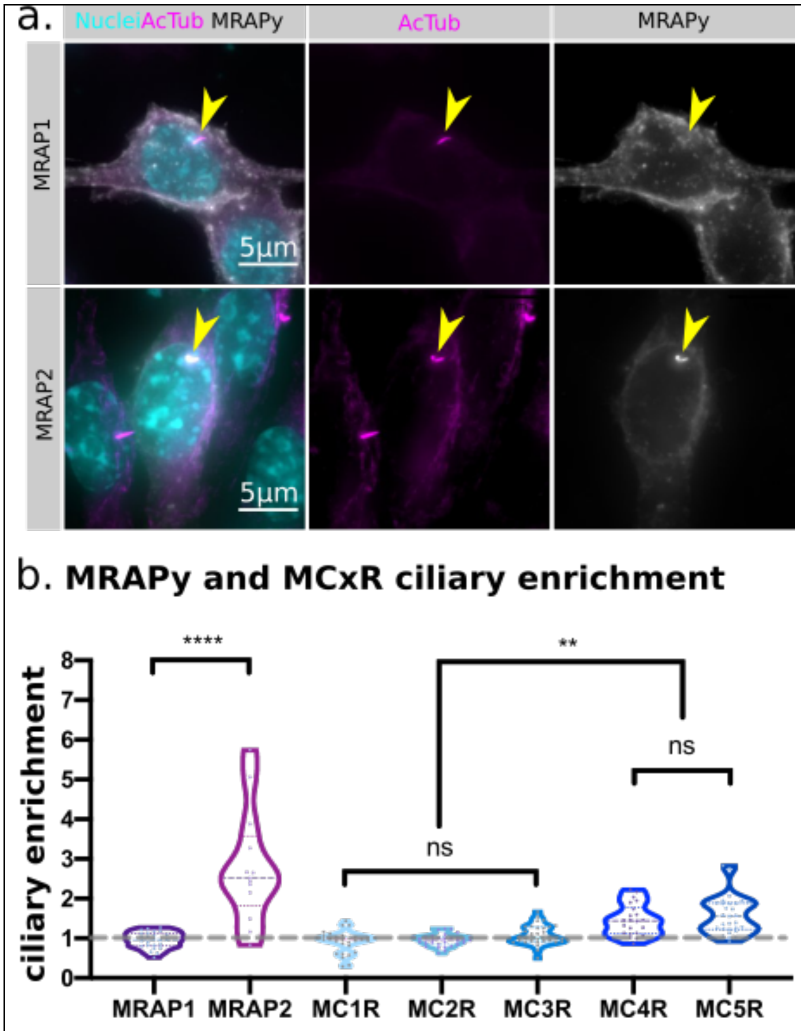


**Figure S2: Ciliary enrichment of Melanocortin receptors, MRAP1 and MRAP2.** ***a*** MRAP2 (lower panel) is enriched at the primary cilium when transfected in IMCD3 cells without any Melanocortin Receptor, while MRAP1 (upper panel) does not. Yellow arrows point at primary cilium of transfected cells. Scale bar 5 µm.***b*** Quantification of the enrichment at the primary cilia of IMCD3 cells transfected with MRAPs and MCRs alone. 30-34 ciliated cells per condition were imaged and analyzed. Ciliary and cell body intensity of MCxR and MRAPy was measured with Fiji. Enrichment at the cilium is expressed as (integrated density at the cilium)/ (integrated density in the cell body). Enrichment >1 indicates higher localization of the protein at the cilium than at the cell body. Data are represented as violin plots, *p<0.05, ***p<0.001, ****p<0.0001, ordinary one-way ANOVA with Sidak's multiple comparisons test.

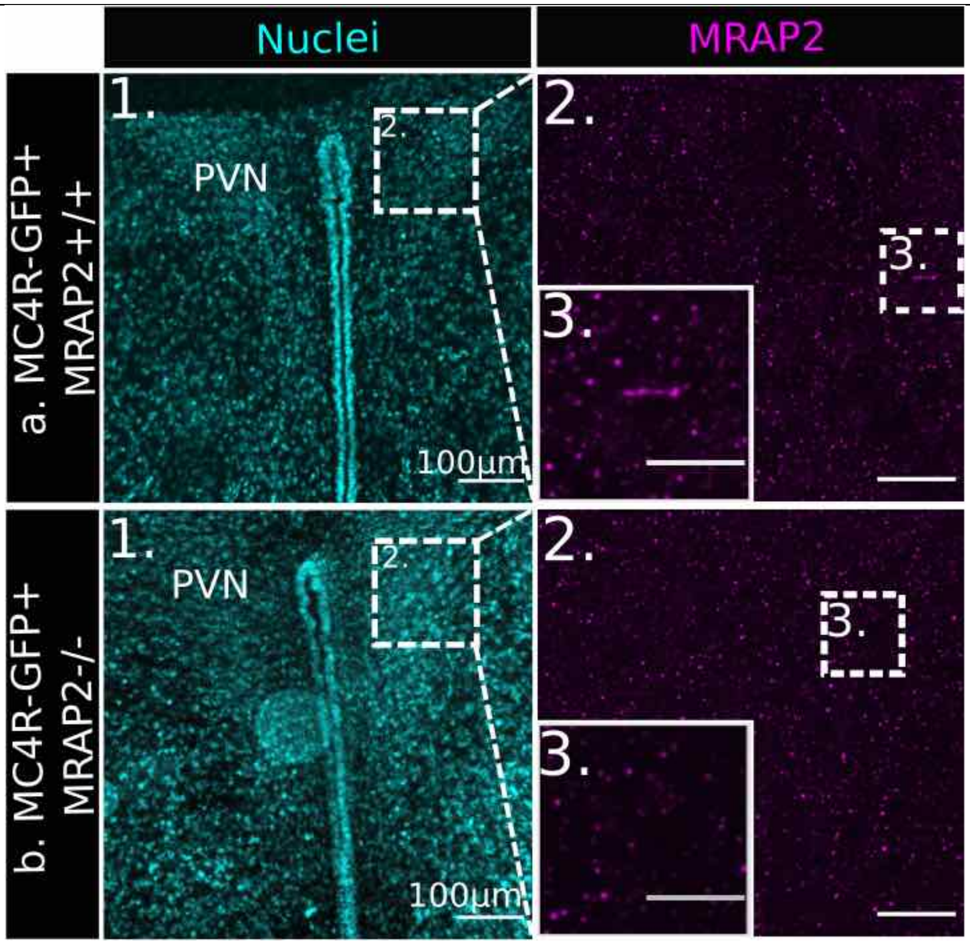


**Figure S3: Specificity of MRAP2 antibody: no signal observed in MRAP2 knock-out hypothalamic sections.** ***a*** In wild-type sections, cilia expressing MRAP2 can be found. ***b*** In *MRAP2* knockout sections, MRAP2 is not detectable. ***1)*** low magnification of the Paraventricular nucleus (PVN), nuclei stained with Hoechst (cyan), scale bar=100μm. ***2)*** Insert from 1), scale bar=20 μm. ***3)*** Insert from 2), scale bar=10μm.

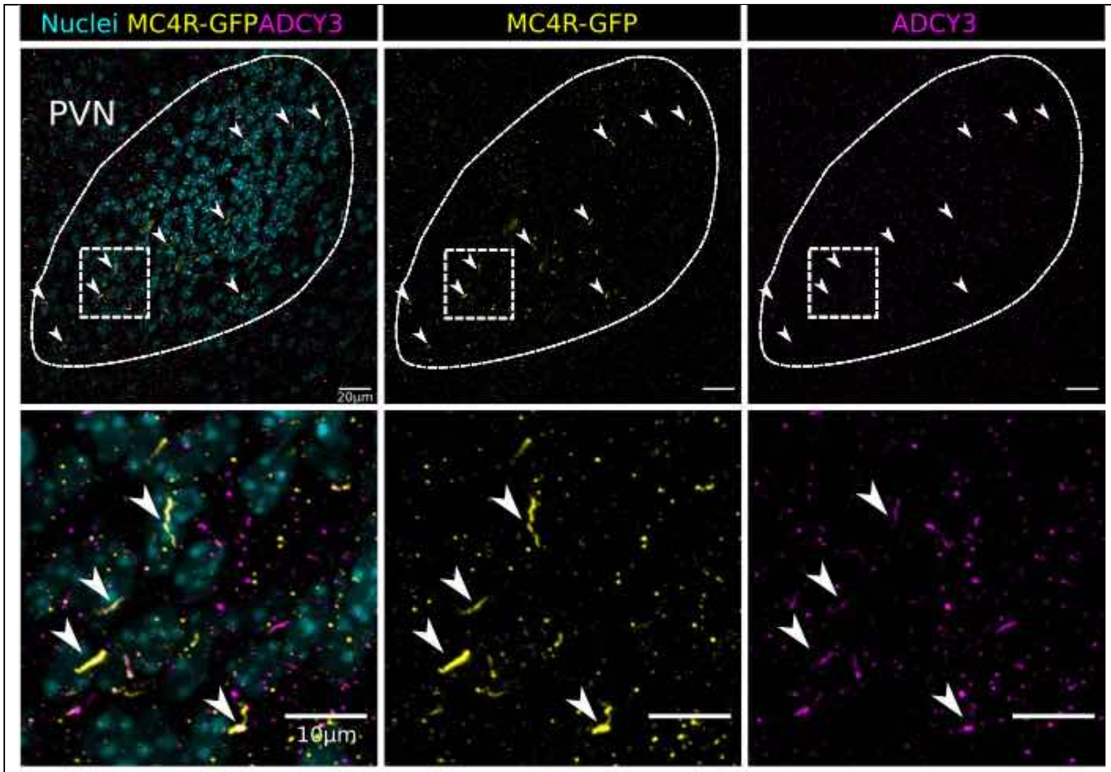


**Figure S4: MC4R localizes at the primary cilium in P6 pups**. MC4R-GFP (Yellow) colocalizes with the specific neuronal primary cilia marker ADCY3 (magenta) in the PVN of P6 pups. Scale bars: 20 μm (top) and 10 μm (bottom).

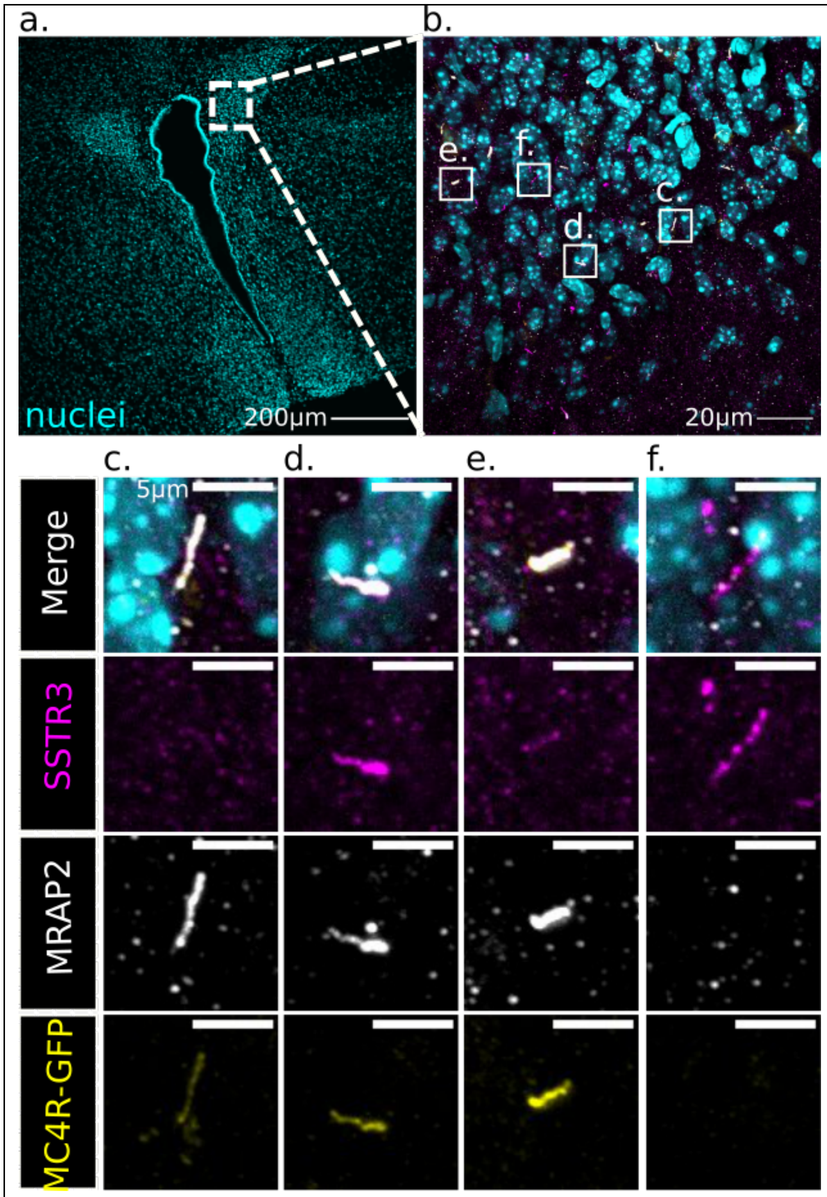


**Figure S5: MRAP2 co-localizes with SSTR3 and MC4R-GFP in a subset of neurons in the PVN**. ***a*** Low magnification image showing the position of the insert in b. Scale bar, 200 μm ***b*** Insert from *a*. Scale bar, 20 μm. ***c*** Primary cilium double positive for MC4R-GFP and MRAP2, but not SSTR3. ***d*** and ***e*** Primary cilia triple positive for MC4R-GFP, MRAP2 and SSTR3. ***f*** Primary cilium positive for SSTR3 only.

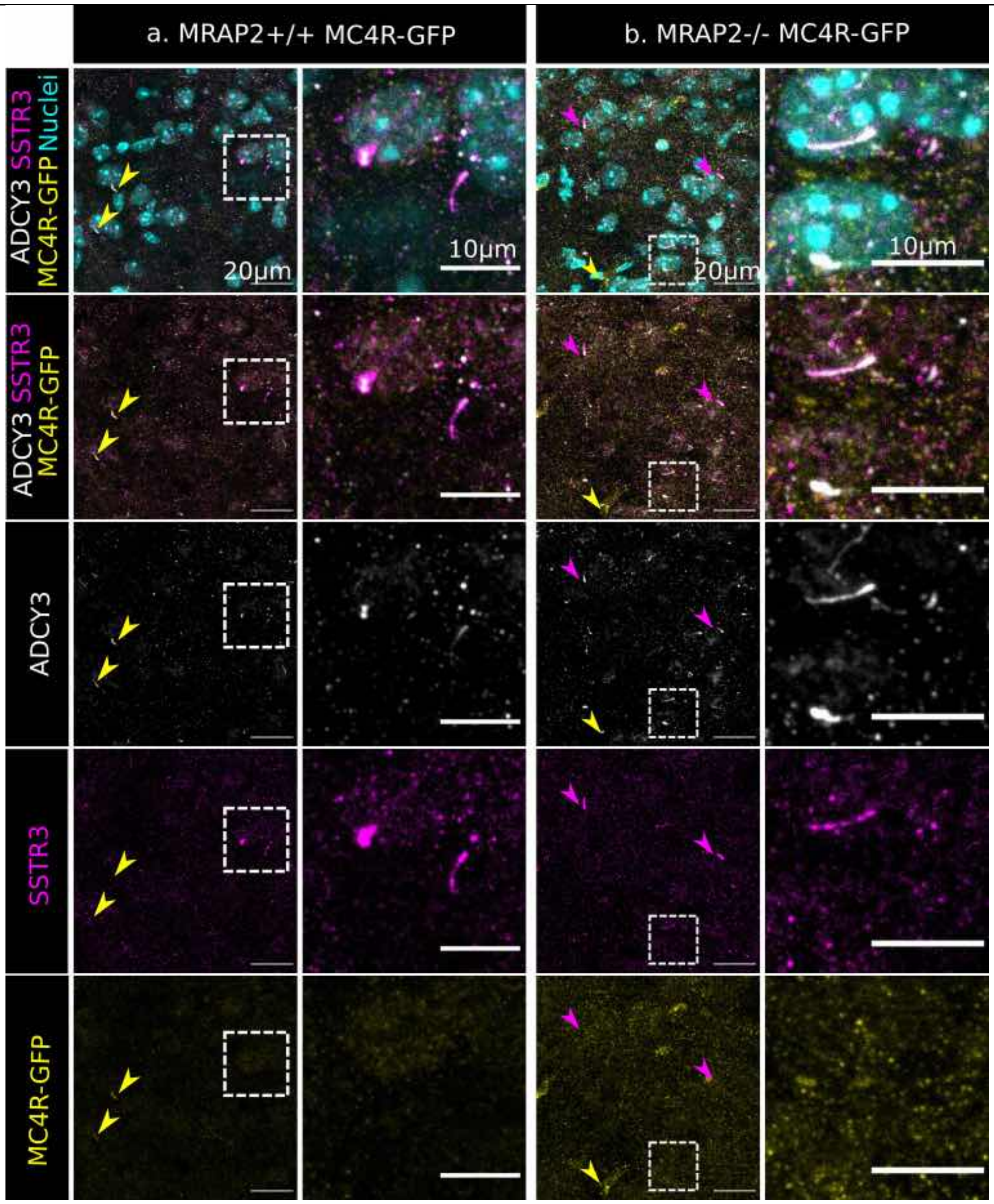


**Figure S6: MRAP2 deletion does not impact SSTR3 ciliary localization.** Immunofluorescence imaging of the PVN of a MRAP2 wild type (***a***) and a MRAP2 knockout mouse (***b***), showing co-localization of SSTR3 (magenta) with ADCY3 (white) at the neuronal primary cilium. Yellow arrows point at MC4R-GFP+ cilia, while magenta arrows point at SSTR3 positive cilia. Second and fourth columns are inserts of first and second columns showing SSTR3+ cilia in both MRAP2 wild type and mutant mice, respectively.
